# FISH-ing for captured contacts: towards reconciling FISH and 3C

**DOI:** 10.1101/081448

**Authors:** Geoff Fudenberg, Maxim Imakaev

**Affiliations:** Center for the 3D Structure and Physics of the Genome, and Institute for Medical Engineering and Science (IMES), Massachusetts Institute of Technology, Cambridge, Massachusetts, USA

**Keywords:** chromosome, polymer, FISH, Hi-C, 3C

## Abstract

Deciphering how the one-dimensional information encoded in a genomic sequence is read out in three-dimensions is a pressing contemporary challenge. Chromosome conformation capture (3C) and fluorescence in-situ hybridization (FISH) are two popular technologies that provide important links between genomic sequence and 3D chromosome organization. However, how to integrate views from 3C, or genome-wide Hi-C, and FISH is far from solved. We first discuss what each of these methods measure by reconsidering available matched experimental data for Hi-C and FISH. Using polymer simulations, we then demonstrate that contact frequency is distinct from average spatial distance. We show this distinction can create a seemingly-paradoxical relationship between 3C and FISH. Finally, we consider how the measurement of specific interactions between chromosomal loci might be differentially affected by the two technologies. Together, our results have implications for future attempts to cross-validate and integrate 3C and FISH, as well as for developing models of chromosomes.

---

> The bewilderments of the eyes are of two kinds, and arise from two causes, either from coming out of the light or from going into the light
>
> — -- Plato, The Allegory of the Cave

## Introduction

While genomes are often considered as one-dimensional sequences, they are also physically organized in three-dimensions inside the cell nucleus, with far-reaching consequences (Dekker and Mirny, 2016; Gibcus and Dekker, 2013; Gorkin et al., 2014). One of the many important implications relates to gene regulation: regions of regulatory DNA are often very far away in the linear genomic sequence from the genes they regulate, yet their regulatory interactions are presumed to rely on direct encounters in three-dimensional space. Ideally, one would be able to follow the exact spatial position of two interacting loci in real time, while simultaneously assaying their functional state and the consequences of their encounters, such as RNA production. However, no method currently exists that allows both directly tracing chromosomes with nucleosome-level spatial resolution, let alone in living cells; current methods are indirect readouts of chromosomal organization. Below we focus on two widely used techniques for assaying chromosomes, DNA-FISH and chromosome conformation capture.

Chromosome conformation capture (3C) techniques have become popular for their high-throughput ability to connect spatial information to the genomic sequence (Bonev and Cavalli, 2016; Dekker et al., 2002; Denker and De Laat, 2015; Schmitt et al., 2016). 3C-based approaches use crosslinking and ligation to capture the information that two genomic loci were spatially proximal. Moreover, 3C techniques are readily generalized to genome-wide scale, usually termed Hi-C (xLieberman-Aiden et al., 2009); here we refer to the contact probability frequency from Hi-C and 3C interchangeably. Importantly, while 3C records whether two loci were in contact in some fraction of cells in the population, it does not record where in the nucleus this contact occurred. Also, 3C is usually performed on large populations of cells. This is advantageous in that 3C can assay both very frequent and very rare events, as the large population allows for a large dynamic range. However, information regarding cell-to-cell variability is not available in population-average maps of chromosomal contact frequencies (Imakaev et al., 2015b; Nagano et al., 2013).

Fluorescence in-situ hybridization (FISH) technologies are appreciated for their ability to specifically determine the spatial position of sets of chromosomal loci by imaging (Fraser et al., 2015). FISH is based on optically labeled probes that hybridize to complementary regions of chromosomes. Importantly, as an imaging-based approach, FISH is intrinsically able to probe cell-to-cell variability and directly record spatial position inside the nucleus. Many different labeling approaches have been considered, including labeling pairs of loci (two-loci FISH) (Sachs et al., 1995), as well larger contiguous regions (Shopland et al., 2006), and even whole chromosome painting (Branco and Pombo, 2006; Tanabe et al., 2002). High-throughput (Shachar et al., 2015) and super-resolution (Beliveau et al., 2015; Boettiger et al., 2016; Fabre et al., 2015) FISH approaches are currently in development. Still, obtaining high-resolution pairwise distance distributions for all pairs of loci, i.e. constructing a pairwise distance map similar to a genome-wide Hi-C contact map, currently remains out of reach.

In studies that primarily rely on 3C-based approaches, FISH is often performed on a subset of loci as a validation. Typically, for loci at increasing genomic separations, their average FISH spatial distance increases and 3C contact frequency decreases (Giorgetti et al., 2014; Hakim et al., 2011; Kalhor et al., 2011; Lieberman-Aiden et al., 2009; Rao et al., 2014). This trend is exactly what is expected for a sufficiently homogeneous polymer, where average spatial distance is perfectly inversely related to contact frequency (Fudenberg and Mirny, 2012; Lua et al., 2004; Rosa et al., 2010). Additionally, for a limited set of tested pairs of loci, it was found that loci in the same A/B compartment contact each other frequently and are on average closer (Lieberman-Aiden et al., 2009), with similar findings for loci in the same TAD (Dixon et al., 2012; Nora et al., 2012). Moreover, spatial distance and contact frequency are largely correlated at the ~300kb-10Mb scale (Wang et al., 2016). However, this does not strictly seem to be the case for all loci (Imakaev et al., 2015b), and can even lead to seemingly-paradoxical observations when comparing FISH and 3C (Williamson et al., 2014).

Here we demonstrate how 3C and FISH generally probe different aspects of spatial chromosome organization. We first illustrate how spatial distance and contact frequency can display a seemingly-paradoxical relationship in currently available experimental data, focusing on the simplest two-locus labeling approach for FISH, as it is most directly comparable to 3C. We then study the connection between contact frequency and average spatial distance in a simple polymer model. Our simulations show that a minimal assumption, introduction of a single dynamic loop between two loci, can break the typical relationship between contact frequency and average spatial distance. In short, our results indicate that the impact of locus-specific chromosome organization can impose seemingly-paradoxical relationships between contact frequency and median spatial distance, as sometimes observed in experimental data.

## Results

### 3C and FISH probe different aspects of spatial organization

To investigate the connection between 3C and FISH we focused on the simplest case of both methods, where each method probes the relationship between a pair of loci. A typical goal of a FISH experiment is measuring the distribution of spatial distances between a pair of loci (PDF, Fig. 1). Results from such experiments are often shown as cumulative probability distributions (CDF, Fig. 1 lower) as these do not require binning or density estimation steps to obtain relatively smooth curves for limited numbers of cells. In contrast with FISH, 3C experiments capture rare contacts that occur when these loci are closer than the capture radius imposed by crosslinking and ligation. Roughly, 3C measures the integral of the spatial distance PDF up to the capture radius, or, equivalently, the value of the CDF at the capture radius. Since such small distances are relatively rare, imaging many cells is certainly a requirement for directly comparing 3C contact probabilities with distances measured by FISH.

**Figure 1.**
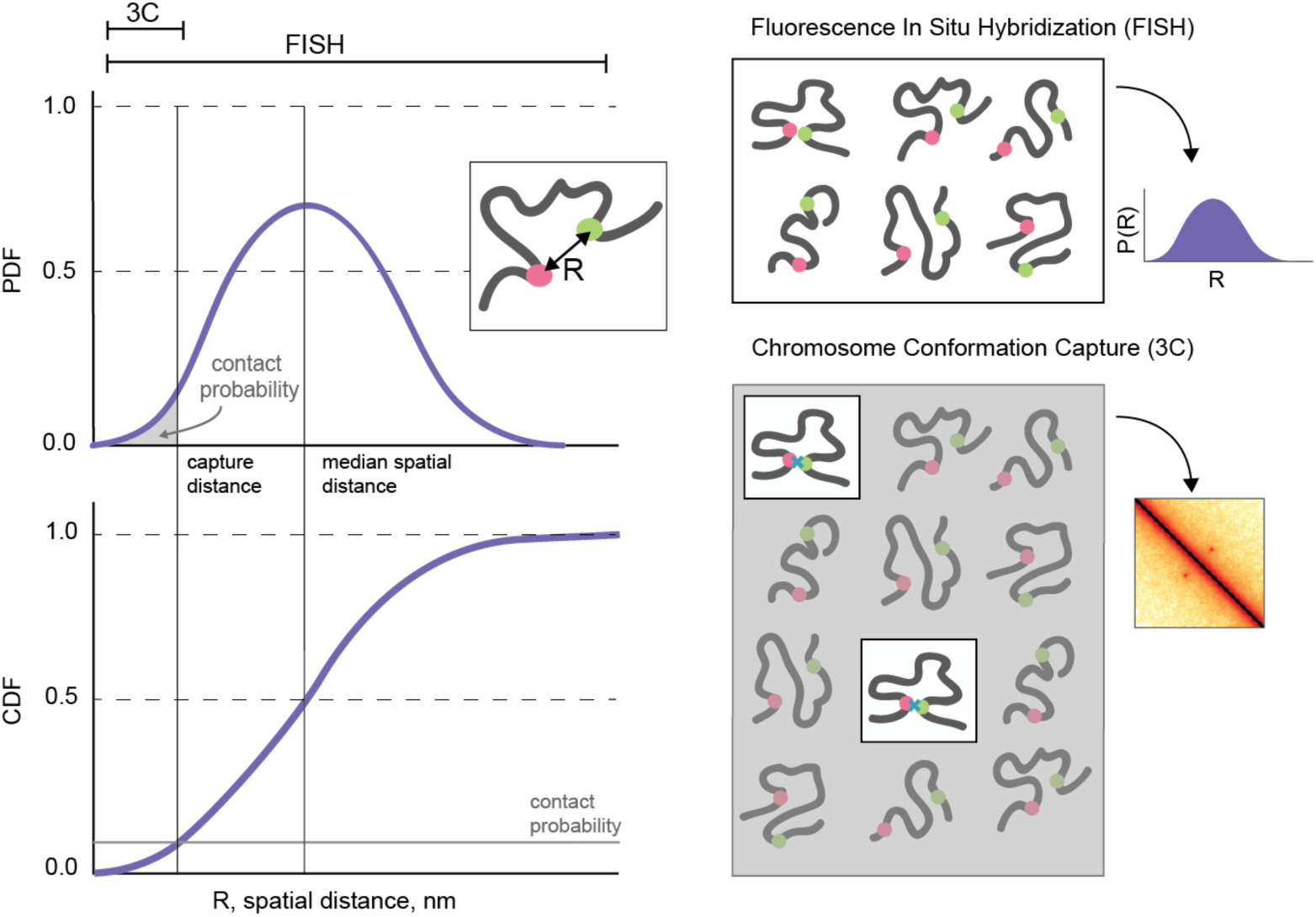
Illustrated relationship between 3C and FISH. (*left*) Illustration of a PDF and CDF pairwise spatial distance, R, between two loci for a large population of cells. Theoretically, FISH can measure the full pairwise spatial distance distribution. 3C captures contacts that occur at distances less than the capture distance, indicated by the area under the PDF up to the capture distance, or the value of the CDF at the capture distance. (*right*) FISH obtains information for all cells in a population to build up a full distribution of pairwise distances between labeled loci. 3C approaches capture contacts from the small fraction of cells where two loci are within the capture radius.

We further examined the connection between 3C and FISH by considering recent publicly-available Hi-C data (Rao et al., 2014).This dataset includes CDF FISH plots for four pairs of ‘loops’ and ‘control’ loci at matched genomic separations, presented as a validation to the Hi-C dataset (GM12878, binned at 10kb, re-plotted in Supplemental Fig 1) for the same cell type. As part of the validation, (Rao et al., 2014) reported that for each of the loop-control pairs of loci, the median spatial distance changed concordantly with the Hi-C signal. While this holds, we also found that this was not always the case when we compared loops and controls from different pairs. Indeed, we found a seemingly-paradoxical relationship between peak-4-loop and peak-3-control; peak-4-loop has roughly 9-fold higher contact frequency despite being further away on average than peak-3¬control. Nevertheless, the change in the value of the CDF at small distances (<300nm) actually was in agreement with Hi-C (Fig 2), suggesting that this short-range behavior of the CDF is more closely connected with contact frequency. A similar situation is observed for peak4-loop and peak-2-control. In contrast, for all control-control pairs of loci, the median spatial distance changed concordantly with the Hi-C signal. Indeed, it is important to note that seemingly-paradoxical pairs involved comparisons with a pair of loop-loci.

**Figure 2.**
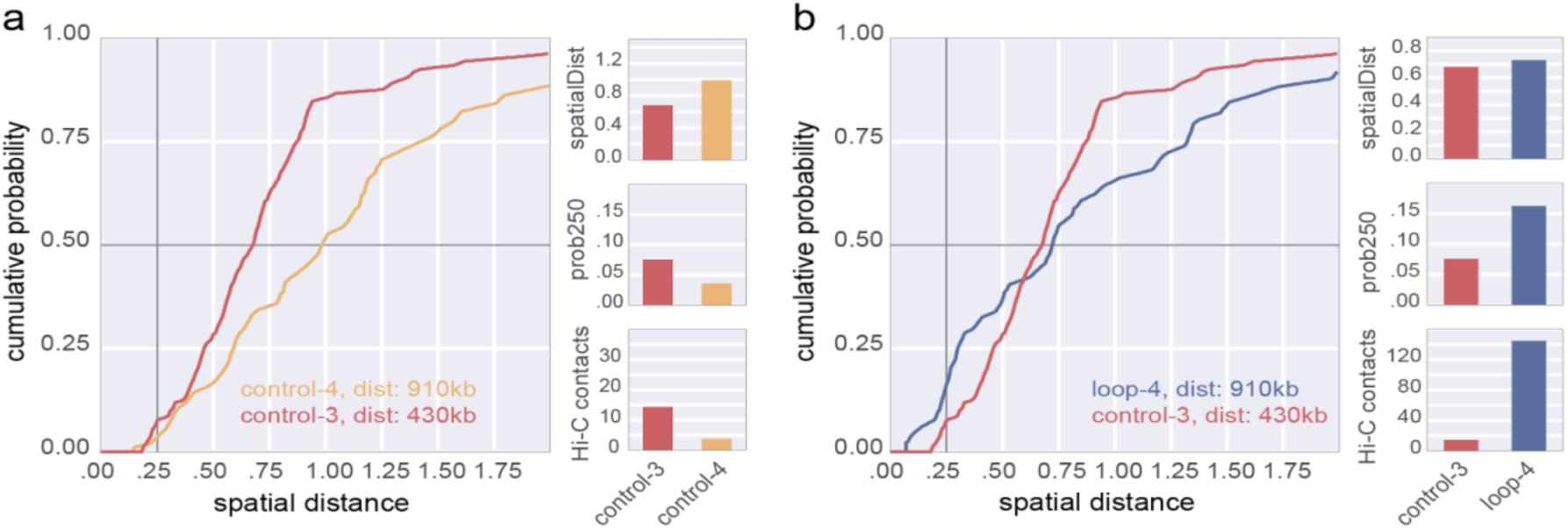
Experimental data demonstrate the complex relationship between Hi-C and FISH. **a**. a comparison between a pair of typical loci shows increased spatial distance, decreased P(dist<250nm), and decreased Hi-C counts. **b.** a comparison between a pair of loop loci and a control pair shows increased spatial distance, but increased P(dist<250nm) and increased Hi-C counts. FISH and Hi-C data replotted from (Rao et al, 2014) for GM12878 cells.

Together, these observations suggest that locus-specific chromosome organization *in vivo* can be an important reason why average spatial distance and contact frequency could behave differently. They additionally suggest that obtaining the full spatial distance distribution, including very short distances, from FISH will be necessary to cross-validate observations from 3C.

### Simulations can reconcile contact frequency and spatial distances

To understand the minimal set of assumptions that can decouple contact frequency from average spatial distance, we investigated both of these quantities in equilibrium polymer simulations of a single dynamic chromatin loop (**Fig 3**). As in our studies of the effect of a fixed chromatin loop (Doyle et al., 2014), we model chromatin as a semi-flexible polymer fiber with excluded volume interactions, 15 nm diameter monomers, each representing roughly three nucleosomes (~500bp), with a persistence length of 3 monomers. To impose excluded volume between non-loop base monomers, we used the repulsive part of the Lennard-Jones potential. To account for the dense arrangement of chromatin within the nucleus (Halverson et al., 2014) we impose a 10% volume density using periodic boundary conditions. We simulated this system with Langevin dynamics using OpenMM (Eastman et al., 2013), and sampled conformations form the resulting equilibrium ensemble. We then calculated simulated spatial distance distributions (Fig 3c,d), contact maps (Fig 3a), and average spatial distance maps (Fig 3b) from these conformations.

**Figure 3.**
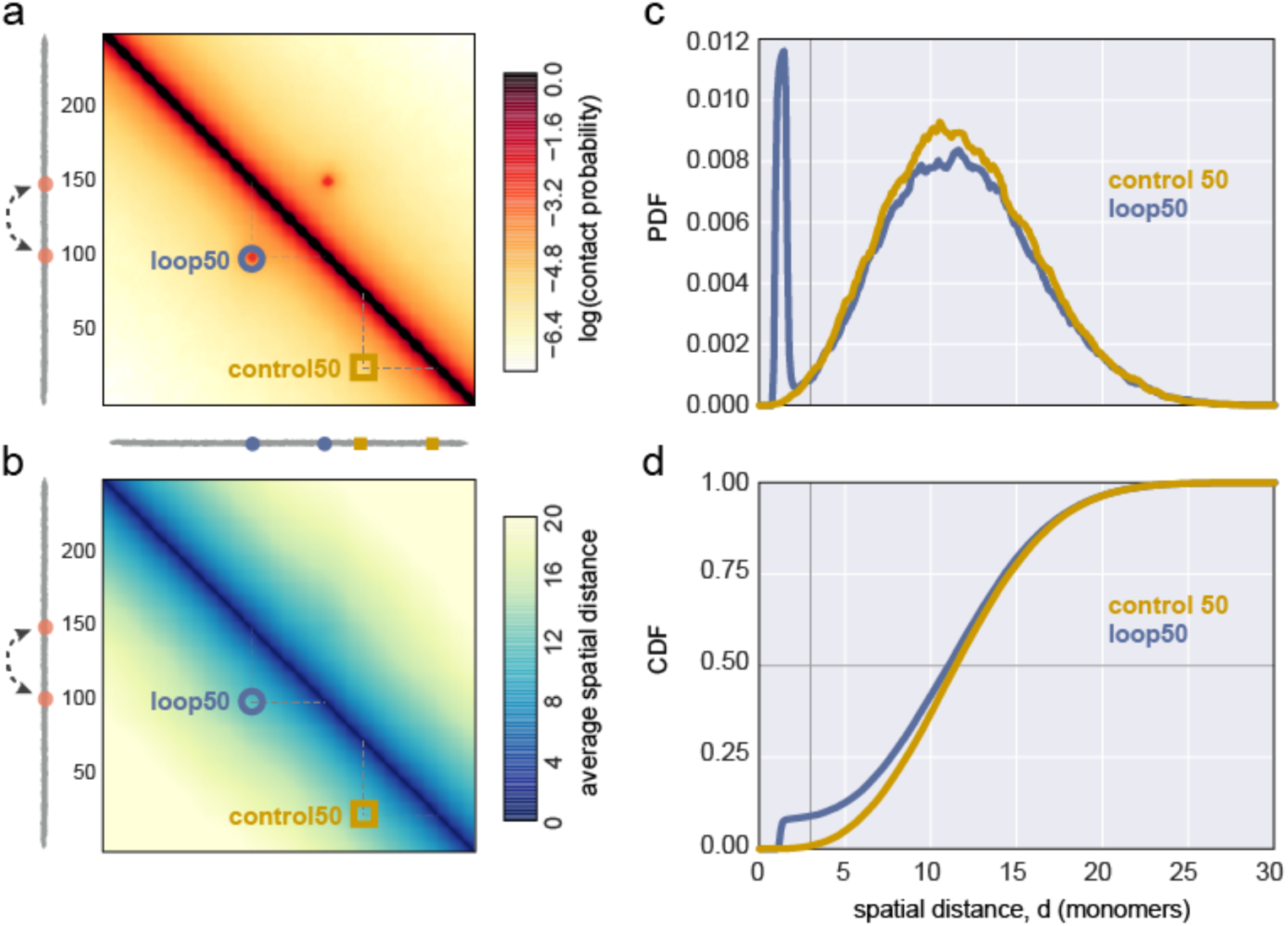
Simulations demonstrate the effect of introducing a single dynamic loop on contact probability maps, and spatial distance distributions. **a.** Contact frequency map for a polymer with a specifically interacting dynamic 50kb loop, indicated as a dashed arc between two loop bases in orange. Two locations for simulated FISH (control50 yellow square, and loop50 blue circle) are indicated on the contact map. control50 indicates an equally spaced pair of control loci, without any specific interactions. **b.** Average spatial distance map for the same region. **c.** PDF of spatial distances for the loop and control pairs of loci (colors as in a). **d.** CDFs for indicated loci; vertical grey line indicates contact frequency, horizontal grey line indicates median distance.

We imposed the dynamic looping interaction using a short-ranged attractive force. Monomers at the base of the 50-monomer dynamic loop interacted with attractive energy (4kT, unless noted) when they were closer than a distance of 2 monomer diameters; for other monomers the attractive part of the potential was set to be negligibly small (0.1kT). This pairwise interaction potential could arise from direct molecular interactions (Barbieri et al., 2012; Jost et al., 2014; Scolari and Lagomarsino, 2015), and can be thought of as imposing a polymer consisting of hard spheres that stick to some degree upon coming into contact (for review, see (Hofmann and Heermann, 2015; Imakaev et al., 2015b)). In our simulation, the dynamic looping interaction is clearly visible in the contact frequency map, but is faint in a map of average spatial distances (Fig 3a,b). Interestingly, the PDF of spatial distances for monomers at the base of the dynamic loop and control monomers (Fig 3c) appeared quite similar, apart from a sharp peak at short distances for the monomers at the loop base. A physical intuition for this behavior comes from considering this result in terms of an equilibrium ensemble of conformations; by introducing a short-range attractive energy, we are simply re-weighting the set of conformations where the loop is present to become more frequent, in proportion to the attraction energy, while other conformations making the bulk of the PDF distribution are uniformly down-weighted.

An important context for our simulations, is that in polymer systems with indistinguishable monomers, the only determinant on both contact frequency and mean spatial distances is the number of intervening monomers between the two loci. For most such polymer systems, at distances much larger than the persistence length, mean spatial distance, *R*, monotonically increases, and contact frequency, *P_*c*_*, monotonically decreases as a function of the number of intervening monomers, *s*. Given a pair of loci, the number of monomers between them, s, fully specifies their contact probability and spatial distance. For example, in a random walk, the distance between monomers, R(s) is proportional to s^1/2^, while the contact probability is inversely proportional to the volume occupied by a section of the polymer, *P*_*c*_*(s)* ~ *R(s)*^*-3*^ = s^-3/2^ Slightly different, yet still directly proportional, relationships can be seen in other polymer ensembles, including the fractal globule and unknotted globules (Bunin and Kardar, 2015; Imakaev et al., 2015a) (to be discussed further in future work). Here, we see this typical, monotonic behavior, in the region of our simulated polymer that lies far from the dynamic loop (Supplemental Fig 2).

However, our simulations demonstrate that even a minimal modification, introduction of a single dynamic loop, changes this typical behavior (Fig 4). While a comparison between control loci of separation 30 or 50 monomers displays the typical monotonic behavior over all genomic separations (Fig 4b), an apparent paradox emerges when comparing the control loci separated by 30 monomers with 50-monomer dynamic loop (Fig 4c). While the 50-monomer dynamic loop is further apart on average, it displays a higher contact probability. This seemingly-paradoxical behavior can emerge in part because contacts are rare events; indeed, because they are rare, contact frequency can increase many-fold without large changes in the average distance (Supplemental Fig 2d,e). This simulation shows how, even in a particularly simple case, seemingly paradoxical relationships can emerge between spatial distance and contact probability, arguing for caution when designing comparisons between FISH and 3C.

**Figure 4.**
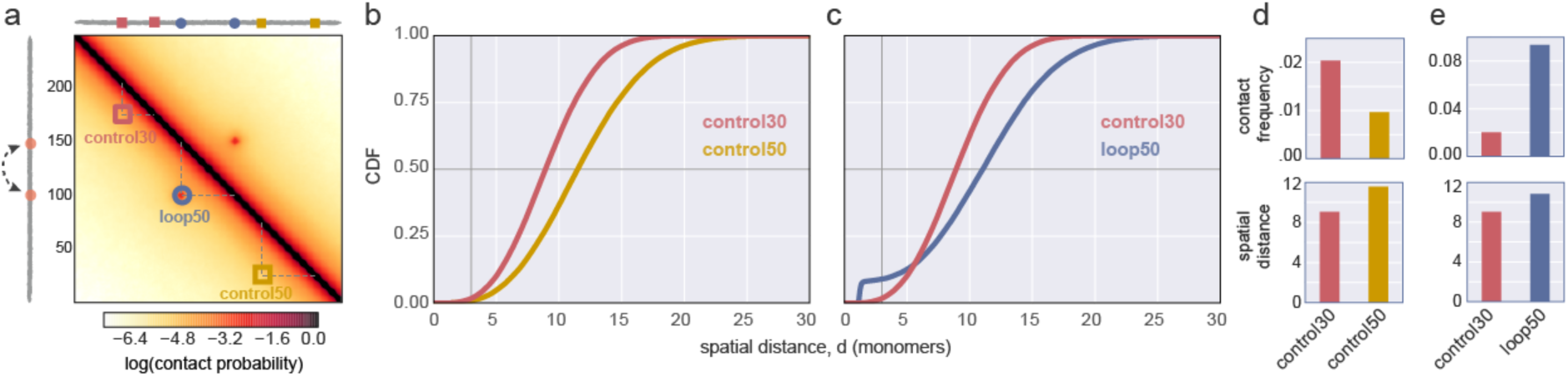
Simulations highlight the complex relationship between contact frequency and median spatial distance. **a.** Contact frequency map for a polymer with a specifically interacting dynamic 50kb loop, as in Fig. 3, indicated as a dashed arc between two loop bases in orange. Three locations for simulated FISH (c30, red square; c50 yellow square, and loop50 blue circle) are indicated on the contact map. c30 and c50 indicate regions without specific interactions between monomers. **b.** Average spatial distance map for the same region. **b,c.** CDFs for indicated loci; vertical grey line indicates contact frequency, i.e. the probability of distance <=3,, horizontal grey line indicates median spatial distance. **d,e.** changes in contact frequency for indicated loci, and median spatial distance for indicated loci pairs. Note that loop50 has a higher contact frequency, but larger median spatial distance, than c30.

### Simulations illustrate how experimental limitations could impact validation of a dynamic loop

We next investigated how possible experimental limitations to either FISH or 3C can impact our ability to ascertain the presence of this simulated looping interaction, as each approach has limitations as compared with the idealized representation in Figure 1 and in polymer simulations.

Crucially for FISH, measuring a finite number of cells imposes uncertainty on the PDF, making the probability of rare events difficult to estimate. Our simulations additionally show that, in general, it is much easier to detect changes in median spatial distance than contact frequency (Supplemental Fig 3). This implies that a consistent validation of Hi-C by FISH, i.e. using only pairs of loci below Hi-C capture distance, would require larger populations of cells than typically used in FISH experiments. For FISH, additional uncertainty can be imposed by factors including: probe size, chromatin movement during hybridization, background noise, and assigning probe pairs to particular chromosomes. In our simulations, we find that even a small uncertainty or imprecision in the spatial localization of probes during FISH makes the existence of a dynamic looping interaction much more difficult to ascertain (Supplemental Fig 4).

For contacts recorded by 3C, the capture radius is imposed by a number of factors, including restriction efficiency, restriction frequency, and the details of crosslinking, which may depend on the particular complement of DNA-associated proteins at a given genomic locus (Belmont, 2014; Gavrilov et al., 2013; Nagano et al., 2015). Indeed, our simulations show that a larger contact radius for simulated 3C can also obscure the existence of a dynamic looping interaction (Supplemental Fig 4). Additional measurement noise may also come from library complexity, sequencing depth, and ligations in solution (Gavrilov et al., 2013; Hsieh et al., 2016; Lajoie et al., 2014; Nagano et al., 2015; Rao et al., 2014), which could all obscure the detection of looping interactions in 3C-based methods.

Together, these simulated perturbations to the idealized FISH and 3C protocols illustrate how considering many experimental details will be required to fully reconcile observations from FISH and Hi-C.

## Discussion

Our results illustrate that while median spatial distance and contact frequency are often inversely proportional, they are far from equivalent. Indeed, our simulations show that a relatively minor assumption-- the existence of a dynamic looping interaction between two loci-- clearly breaks the equivalence between these two quantities. In particular, we show that there is great freedom to make large shifts in contact frequency with small shifts to median spatial distance, since contacts between distal chromosomal loci are generally rare events. This has important implications in vivo, where Hi-C contact maps and FISH reveal many structural elements, including large-scale compartments, TADs, and CTCF-mediated looping-interactions, that can impose even more complex relationships than a single dynamic loop.

Given these factors, 3C experiments cannot be simply validated (or invalidated) by FISH, without carefully considering technical details of the two methods. Indeed, efforts to integrate results from these technologies will need to carefully address unknowns of the 3C capture radius and FISH localization uncertainty, in addition to assaying sufficiently large numbers of cells to populate the small distance portion of the FISH distribution.

In addition to the implications for validation, our results also caution against certain modeling approaches. In particular, our results show how a common strategy (Hu et al., 2013; Lesne et al., 2014; Rousseau et al., 2011; Segal et al., 2014; Trieu and Cheng, 2015; Varoquaux et al., 2014; Zhang et al., 2013) of simply transforming 3C contacts into spatial distances ultimately leads to inconsistent models of chromosomal organization (for review see (Imakaev et al., 2015b)).

Nevertheless, our results also demonstrate how polymer modeling can in principle be used to reconcile FISH spatial distances and 3C/Hi-C contact frequencies. This is because both quantities are readily calculable from an ensemble of simulated polymer conformations. As high-throughput and high-resolution imaging data becomes available, systematically comparing polymer models to both Hi-C and imaging data will be an essential step towards understanding principles and mechanisms of chromosomal organization.

While we limit ourselves to considering the relatively simple comparison between 3C and two-loci FISH, many other comparisons would be valuable in future work. In particular, it is less clear how to best compare information obtained from contiguously stained regions (Boettiger et al., 2016; Shopland et al., 2006) with 3C experiments, as the former constitutes an intrinsically multipoint interaction. Another useful direction would be to consider assays that probe spatial proximity to subnuclear bodies (Mao et al., 2011), including the lamina (Kind et al., 2013, 2015), the nucleolus (Nemeth et al., 2010), and nuclear speckles (Spector and Lamond, 2011. Non-equilibrium effects on chromosomal organization, both following decondensation (Muller et al., 2010; Rosa and Everaers, 2008; Walter et al., 2003), and through interphase (Bancaud et al., 2009; Fudenberg et al., 2015), may also have interesting differential effects on contact frequency and median spatial distance between loci. Methods for time-resolved imaging of chromosomal loci in living cells (Bronstein et al., 2009; Chen et al., 2013; Chuang et al., 2006; Hajjoul et al., 2013; Javer et al., 2014; Weber et al., 2010) are promising approaches for probing these non-equilibrium phenomena.

Finally, the difference between spatial distances and contact probabilities has implications for experimental design in functional studies of gene regulation. Indeed, both gene expression and chromosomal organization are intrinsically stochastic processes. Gene expression occurs in bursts, and is often variable both in time and between cells (Paulsson, 2004; Raj and van Oudenaarden, 2008). Similarly, chromosomal folding is highly variable, and chromosomes adopt a highly diverse set of conformations (Gibcus and Dekker, 2013; Imakaev et al., 2015b; Nagano et al., 2013). Despite this, static depictions of an enhancer-promoter loop are commonly used to illustrate the connection between these two intrinsically stochastic processes. Instead, probabilistic contacts between an enhancer and a promoter form a natural connection between the two intrinsically stochastic processes of gene expression and chromosome folding (Krijger and De Laat, 2013). As such, we propose that experiments probing changes in enhancer-promoter contacts, rather than average enhancer-promoter distance, will have a much better chance of unraveling causal relationships between chromosomal organization and gene expression.

## Supplemental Figures and Methods

**Supplemental Figure 1.**
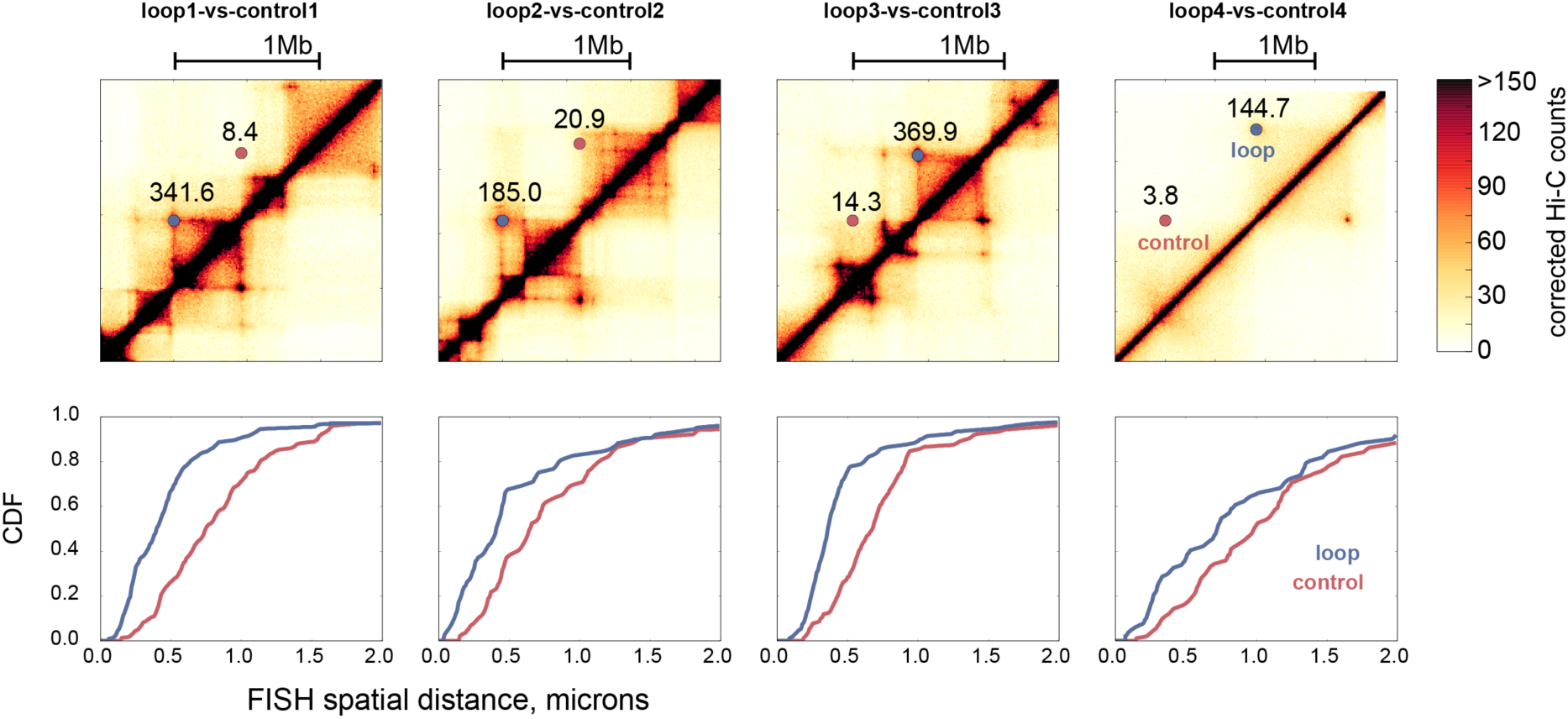
FISH and Hi-C data reproduced from Rao et al., 2014. *Top*: Hi-C maps for the probed regions, numbers indicating the corrected Hi-C counts at indicated loop and control loci. *Bottom*: CDFs for FISH, loop loci in blue, equal genomic separation control loci in red.

**Supplemental Figure 2.**
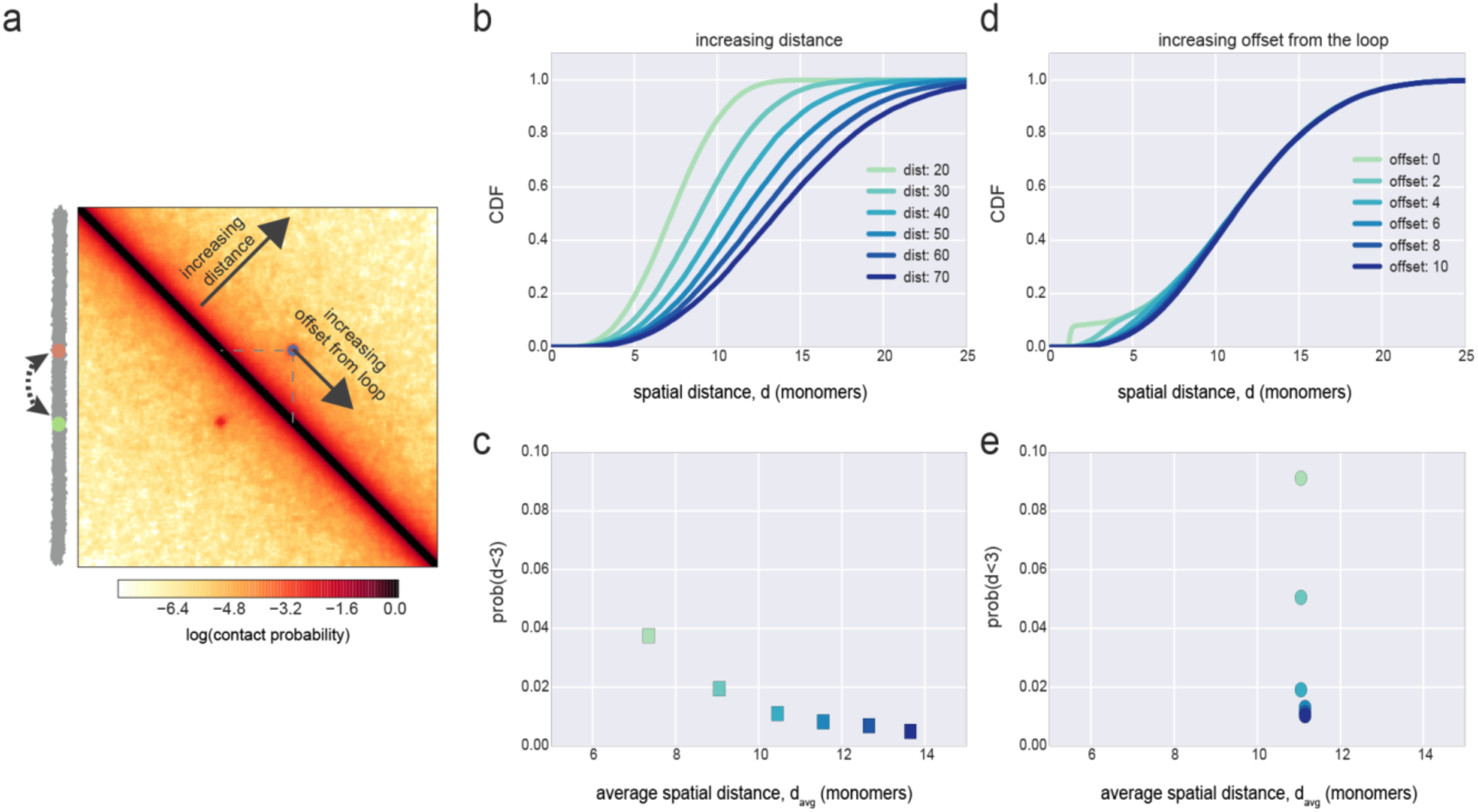
**a.** Average contact map, as in main Fig 2. Arrows show position of probes for increasing distance away from the dynamic loop, and for increasing offset from the loop at a fixed distance. **b.** CDFs for loci pairs at increasing genomic separations. Note that these loci-pairs are chosen to be away from the dynamic loop. **c.** Relationship between average spatial distance and contact frequency (distance < 3 monomers) for these loci. **d.** CDFs for loci pairs at increasing offsets from the bases of the dynamic loop, but the same genomic separation. **e.** Relationship between average spatial distance and contact frequency (distance < 3 monomers) for these loci. Note that average spatial distance clearly increases as contact frequency decreases in the first case, but not in the second. Indeed, as **d** and **e** show, contact frequency can increase many-fold without large changes in the average distance

**Supplemental Figure 3.**
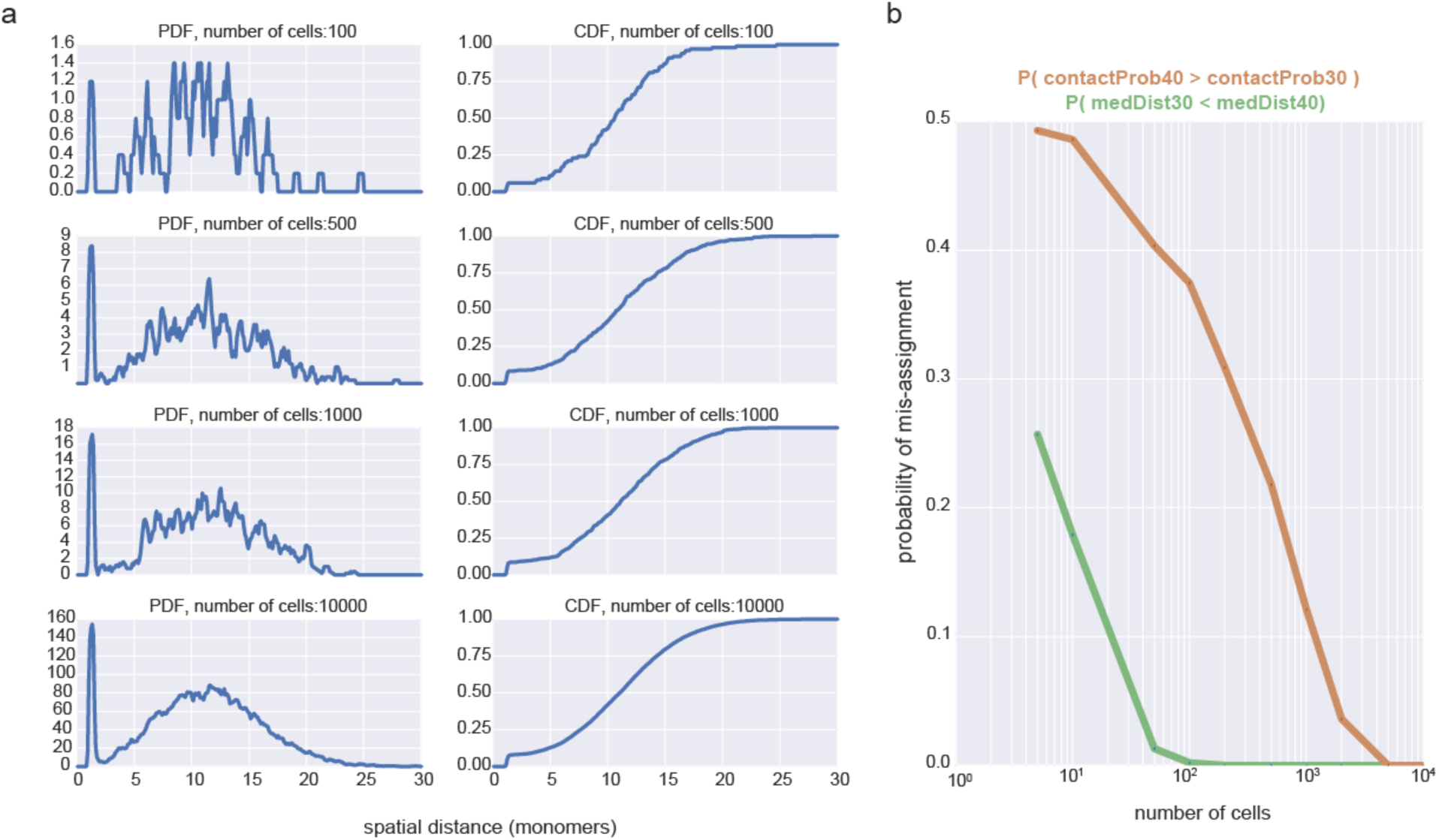
**a.** Simulated PDFs and CDFs for increasing number of sampled conformations the two loci at the base of a simulated dynamic loop. **b.** probability of observing the incorrect relationship between median distances (green) or contact probabilities (orange) for two loci separated by either 30 or 40 monomers. These curves illustrate how many more cells are needed to reliably estimate contact frequency than median spatial distance. Strikingly, the 2-fold difference in contact frequency is much more difficult to reliably detect than a 1.17-fold difference in average distance with a limited number of cells.

**Supplemental Figure 4.**
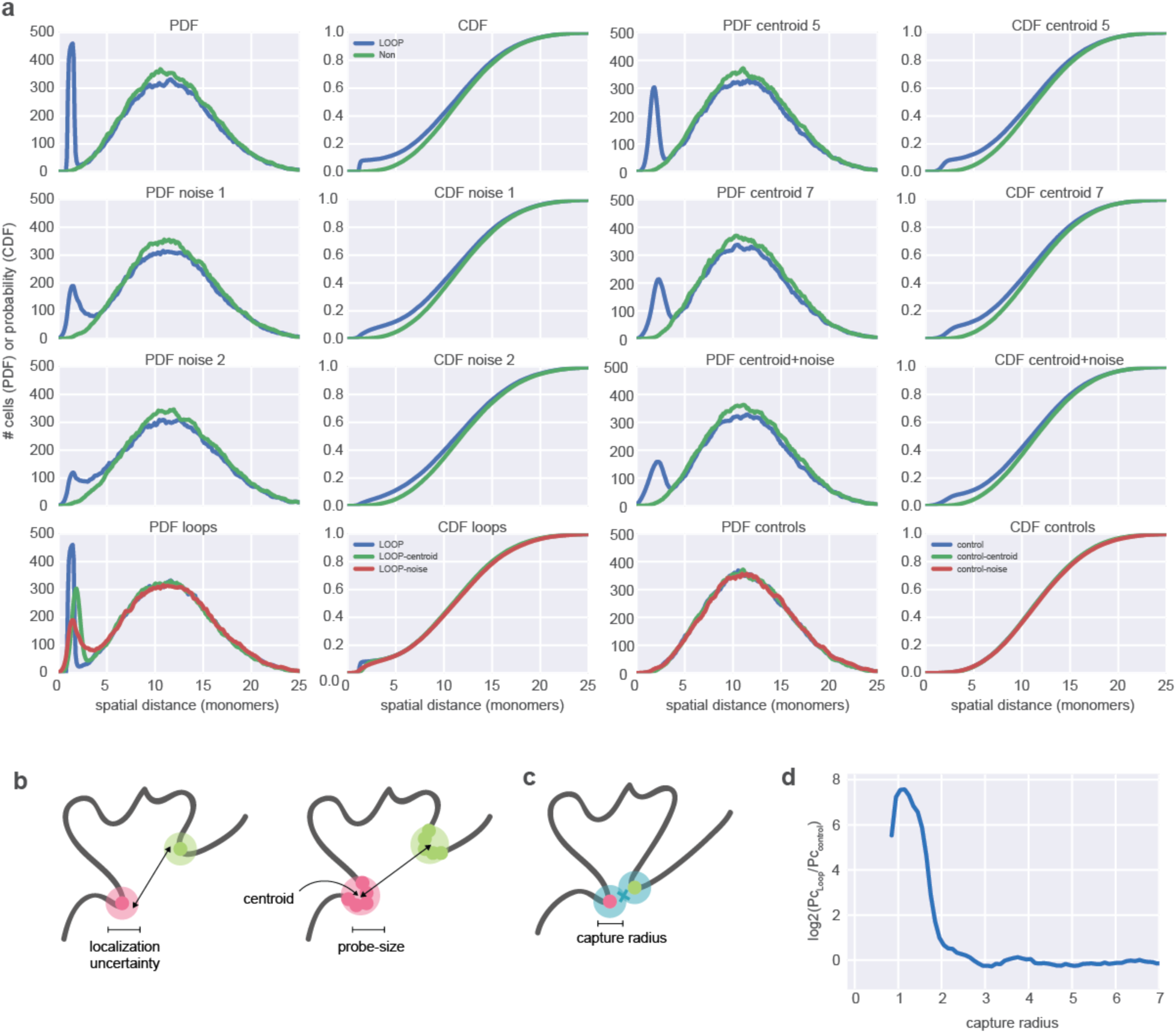
***a.*** The effect of localization uncertainty or size of simulated FISH probes on measured PDF and CDF. Subplots show the indicated PDF or CDF pairs for the loop region or the control region at the same monomer separation calculated over 39665 conformations. Probe localization uncertainty is simulated by adding Gaussian noise with the indicated width to each simulated set of probe distances. Probe size is simulated by considering the distance to the centroid of a labeled region of the indicated size. Note that even a 2-monomer uncertainty in localization can drastically diminish the visibility of a peak, much more so than measuring centroid-to-centroid distances of a pair of 7-monomer regions instead of two 1-monomer probes. We also note that (Giorgetti et al., 2014) found a 35nm uncertainty led to best agreement between their polymer models and FISH data, which would correspond to a 3.5 monomer localization uncertainty in the scenario we present. Also note that the control distribution changes very little for any of these perturbations. ***b.*** Illustration of localization uncertainty (left) or size (right) for FISH probes; size of a probe imposes a centroid-to-centroid measurement, averaged over the positions of all monomers labeled by the probe. **c.** Illustration of capture radius for 3C. ***d.*** The log2(ratio) of contact frequency for the 50 monomer loop considered in the main text divided by the non-loop control at the same distance, as a function of capture distance (in simulated 3C). Note that this decreases to zero after around 3 monomers.

## Methods

### Polymer Simulations

As previously (Doyle et al., 2014; Naumova et al., 2013), we modeled chromatin as a fiber of monomers. Unless noted, each spherical monomer had a diameter of 10 nm and represented 500bp, or approximately three nucleosomes. Adjacent monomers were connected by harmonic bonds with a potential U = 25*(r – 1)^2^ (here and below, energy is in units of kT). The stiffness of the fiber was modeled by a three point interaction term, with the potential U = k*(1-cos(α)), where α is an angle between neighboring bonds, and *k* is a parameter controlling stiffness, here set to 2kT. To model the dynamic loop considered in the present work, we used a Lennard-Jones (LJ) potential U = 4ε_ij_ * (1/r^12^ - 1/r^6^) where ε_ij_ was set to 4kT for the monomers at the base of the dynamic loop (i,j), and was set to negligibly small otherwise (ε= 0.1kT).

Polymer models were simulated with OpenMM (Eastman et al., 2013), a high-performance GPU-assisted molecular dynamics software (https://simtk.org/home/openmm). We used an in-house *openmm-polymer* library (publicly available http://bitbucket.org/mirnylab/openmm-polymer). We initialized our simulations as a system of 8 compact rings (see (Imakaev et al., 2015a)), and used periodic boundary conditions to achieve a density of 0.10. We then simulated 50 runs of this system using Langevin Dynamics, for 10e8 time steps. For the fiber lengths considered here, polymer simulations reached equilibrium in less than 1e7 time steps; this was confirmed by observing that monomer displacement saturates after about 5e6 blocks. Conformations have been stored every 1e5 time steps and an equilibrium ensemble of 900 conformations obtained after the initial equilibration was used for our analysis. An Andersen thermostat was used to keep the kinetic energy of the system from diverging using a time step that ensured conservation of kinetic energy.

To obtain simulated contact maps, we first found all contacts within each polymer conformation, and then aggregated these contacts for all pairs of monomers. A contact was defined as two monomers being at a distance less than Rc=3 monomer diameters. To obtain simulated FISH distributions, we calculated a list of spatial distances for a chosen set of loci, and built a histogram of distances starting at 0 in bins of 0.1 monomers. To display PDFs this histogram was then smoothed with a moving average window with a size of 0.7 monomers.

### Experimental data

FISH CDFs were obtained digitized coordinates from published data (Rao, 2014) using webplotdigitizer http://arohatgi.info/WebPlotDigitizer/app/ in the automatic mode with dx=.01 and x-step-with-interpolation. Published Hi-C data (Rao, 2014) was re-processed, filtered, and iteratively corrected using hiclib http://mirnylab.bitbucket.org/hiclib/ (Imakaev et al., 2012).

## Acknowledgements

The authors thank Anton Goloborodko for thoughtful comments and the statistical mechanics analogy of re-weighting loop conformations, and Guy Nir for helpful discussions regarding imaging. The authors also thank Job Dekker and other members of the UMass-MIT Center for 3D Structure and Physics of the Genome for helpful feedback. Finally, the authors thank Leonid Mirny for comments on earlier drafts of this paper, and for supporting their independent. This work was supported by NSF 1504942 Physics of Chromosomes (PI: Leonid Mirny) and U54 DK107980 3D Structure and Physics of the Genome (PIs: Job Dekker and Leonid Mirny).

